# Ghost interactions in MEG/EEG source space: A note of caution on inter-areal coupling measures

**DOI:** 10.1101/220459

**Authors:** J. Matias Palva, Sheng H. Wang, Satu Palva, Alexander Zhigalov, Simo Monto, Matthew J. Brookes, Jan-Mathijs Schoffelen, Karim Jerbi

**Author notes:** Correspondence should be addressed to J. Matias Palva, Neuroscience Center, University of Helsinki, Helsinki, Finland.

## Abstract

When combined with source modeling, magneto‐ (MEG) and electroencephalography (EEG) can be used to study long-range interactions among cortical processes non-invasively. Estimation of such inter-areal connectivity is nevertheless hindered by instantaneous field spread and volume conduction, which artificially introduce linear correlations and impair source separability in cortical current estimates. To overcome the inflating effects of linear source mixing inherent to standard interaction measures, alternative phase‐ and amplitude-correlation based connectivity measures, such as imaginary coherence and orthogonalized amplitude correlation have been proposed. Being by definition insensitive to zero-lag correlations, these techniques have become increasingly popular in the identification of correlations that cannot be attributed to field spread or volume conduction. We show here, however, that while these measures are immune to the direct effects of linear mixing, they may still reveal large numbers of spurious false positive connections through field spread in the vicinity of true interactions. This fundamental problem affects both region-of-interest-based analyses and all-to-all connectome mappings. Most importantly, beyond defining and illustrating the problem of spurious, or “ghost” interactions, we provide a rigorous quantification of this effect through extensive simulations. Additionally, we further show that signal mixing also significantly limits the separability of neuronal phase and amplitude correlations. We conclude that spurious correlations must be carefully considered in connectivity analyses in MEG/EEG source space even when using measures that are immune to zero-lag correlations.

**Highlights:**

✓ Reliable estimation of neuronal coupling with MEG and EEG is challenged by signal mixing
✓ A number of coupling techniques attempt to overcome this limitation by excluding zero-lag interactions
✓ Contrary to what is commonly admitted, our simulations illustrate that such interaction metrics will still yield false positives
✓ Spurious, or “ghost”, interactions are generally detected between sources in the vicinity of true phase-lagged interacting sources
✓ Signal mixing also severely affects the mutual separability of phase and amplitude correlations

## 1 Introduction

Inter-areal interactions among neuronal ensembles during rest or in active tasks are a hallmark of integrative brain function and have been the focus of a thriving body of research over the last decade [Bastos and Schoffelen, 2016; Biswal et al., 2010; Brookes et al., 2011; Foster et al., 2016; Harris and J. A. Gordon, 2015; Hutchison et al., 2013; Karl J., 2011; Mantini et al., 2007; Pizzella et al., 2014; Schoffelen and J. Gross, 2009; Siems et al., 2016; Sporns, 2015; van Diessen et al., 2015]. Magneto‐ (MEG) and electro-encephalography (EEG) offer a highly valuable approach for probing these interactions both by yielding direct electrophysiological recordings of neuronal activity, whole-head coverage and, most importantly, the millisecond-range temporal resolution required for observing fast neuronal dynamics. However, limited spatial resolution and signal processing complexities require attention to subtleties in the obtained coupling results and may lead to erroneous interpretations of the data.

A central problematic issue results from signal spread, which translates to volume conduction in the case of EEG recordings, to field spread when it comes to MEG, and to signal leakage in source reconstructed EEG or MEG data. In both MEG and EEG, a spatially widespread group of sensors detects the activity of any single neuronal source. Therefore, correlations among signals measured at two distant sensors do not necessarily reflect the existence of two distinct interacting cortical sources. On the other hand, from the perspective of individual sensors, the same sensor can always pick up multiple sources. Thus, two instantaneously interacting (*i.e.*, zero phase lag) sources are difficult to be distinguished from a single source whose activity recorded by the same sensors. In addition to these theoretical limitations due to signal spread effects, difficulties in relating results of sensor-level interaction analyses to known anatomical or functional systems, even if caused by true interactions, provide further arguments to why in general interaction analyses should not be performed in sensor space.

The application of source estimation techniques to MEG/EEG data, followed by performing interaction analyses on reconstructed source activations, alleviates but does not fully solve the detrimental effects of signal spread [Gross et al., 2013; Palva and J. M. Palva, 2012; Schoffelen and J. Gross, 2009]. Inverse modeling techniques use spatiotemporal channel information and provide a plausible distribution of neuronal currents that may have generated the sensor-level measurements. The properties (*e.g.*, the spatial smoothness) of the reconstructed source activity depend on the assumptions on which the inverse operator is built and vary across different inverse solutions [Baillet and Garnero, 1997; Gross et al., 2001; Hamalainen and R. J. Ilmoniemi, 1994; Van Veen et al., 1997]. No inverse solution, however, is perfect, and the interpretation of analysis results based on source reconstructed data should always consider the inherent spatial limitations of the inverse technique used. *I.e.*, residual signal leakage will always characterize the source data. Generically, these spatial limitations can be investigated using realistic simulations that employ accurate forward models, in order to evaluate the inverse technique’s point spread (PSF) and cross-talk functions (CTF) [Hauk and Stenroos, 2014; Hauk et al., 2011; Korhonen et al., 2014; Liu et al., 2002; Lütkenhöner, 2003]. These functions quantify, as a function of space, for any given source location, the extent to which the activity at the given location leaks to other locations (PSF), and the extent to which activity that leaks from other locations affects the estimate of the source activity at the given location (CTF). Both measures can be obtained from the so-called resolution matrix, which is the product of the inverse and forward operator matrices.

The detrimental effect of spatial imperfections in the inverse operator manifests itself clearly in the context of interaction analyses between estimated source time courses. Conceptually, the estimated interactions can be driven either by (a) true, (b) artificial or (c) spurious interactions among the reconstructed signals. These notions are defined in this study as follows:

*True* interactions: these reflect estimated interactions that are caused by real interactions between neuronal groups observed at the considered locations.

*Artificial* interactions: these reflect estimated interactions that are false positives and not caused by real interactions between neuronal groups at the considered locations. Rather, the ‘significant’ coupling is caused by signal mixing and often through cross-talk from dominant sources at other locations and thus reflects residual effects of the signal spread at the source level. One well-known example of this is sometimes referred to as ‘seed blur’.

*Spurious* interactions: these reflect estimated interactions that are false positives and also result from cross-talk [Palva and J. M. Palva, 2012]. Yet, the distinction with the artificial interactions described above is that the process underlying the estimated interaction is a genuine interaction between neuronal groups but the location of the interacting sources is misestimated. Concretely, signal spread results in pairs of sources in the vicinity of the actual interacting sources to also display significant coupling. In other words, spurious interactions arise as an unwanted by-product of a truly interacting pair of sources, and can be referred to as *ghost interactions*.

One commonly used strategy to minimize false positives in interactions estimated with MEG is to use an experimental or baseline contrast in combination with either standard (*e.g.*, coherence, amplitude correlations etc.) or signal-mixing insensitive (as described below) interaction measures, and assume that the spatial structure in the false positives is similar across conditions. Obviously, this strategy is only applicable in situations where such contrasts can be made, and therefore it is not applicable in task-free (resting state) situations. More importantly, the validity of the interpretations heavily relies on the untenable assumption that the false positives are similar across conditions. For instance, differences in signal-to-noise ratio result in trivial differences in false positive differences in interactions [Bastos and Schoffelen, 2016].

Recent years have witnessed the development of important and innovative measures that directly avoid false positive observations of coupling attributable to signal spread [Brookes et al., 2012; Hipp et al., 2012; Nolte et al., 2004; Vinck et al., 2011]. These methods exclude the contribution of instantaneous signal spread to the estimated interactions, and by design, thus address the issue of artificial interactions. For instance, the imaginary part of coherency [Nolte et al., 2004] removes the zero-phase lag interactions because these are entirely captured by the real part of coherency. Another types of measures aim to quantify the correlation in band-limited amplitude envelopes. Here, the signals are orthogonalized with respect to each other to remove zero-lag mixing prior to computing the correlation between the amplitudes [Brookes et al., 2012; Hipp et al., 2012].

While these methods can be very useful, they have an important limitation. Ignoring near-zero-lag interaction components makes the interaction estimate insensitive to leakage, also true near-zero-phase interactions will remain undetected.

One important and frequently overlooked limitation of the above-mentioned leakage insensitive measures of interactions is that these measures do not protect against false positives due to spurious interactions, as defined above. Steps towards addressing this problem have already been taken in the case of amplitude correlations [Colclough et al., 2015], but generic interaction-metric independent solutions have remained elusive. Another subtle but equally important problem is the fact that, due to the unavoidable signal leakage, orthogonalized envelope correlation estimates may be affected by the concurrent presence of phase coupling. These limitations pose important challenges to the physiological interpretability of the results. Although the issue of spurious interactions has been recognized by some experts in the field, it and the possible confusion of phase and amplitude couplings are not common knowledge and hence merit more widespread awareness. The purpose of this study is to demonstrate and quantify these ghost interactions and further elucidate the effects of phase coupling on orthogonalized amplitude correlation estimates.

**Footnote**: the term “artificial” and “spurious” interactions are often used interchangeably in the literature. Here, spurious (or ghost) interactions refers only to false positives that arise independently of the chose interaction metric. Spurious/ghost interactions in this meaning have also been termed “inherited” interactions [Hauk and Stenroos, 2014; Colclough et al., 2015]. Furthermore, we do not discuss higher order artificial interactions, i.e. caused by common drive, third-party sources and cascade effects, although identifying them is of equal importance [Mannino and Steven L. Bressler, 2015; Mannino and Steven L. Bressler, 2015; Wollstadt et al., 2015].

## 2 Materials and Methods

### 2.1 Simulation of signals and interactions

‘Estimated source signals’ were modelled as an instantaneous linear mixture (to model signal spread) of underlying source time series. To model these time series, we applied a two-stages mixing procedure.

At the first stage, we modelled the underlying ‘true’ source time series as follows: One-dimensional random Gaussian time series *n*_*i*_ were linearly mixed using mixing parameters *C*_*A*_ and *C*_*θ*_. The mixed time series were filtered using complex Morlet wavelets, and time series to be used as instantaneous amplitudes and phases were computed as follows,

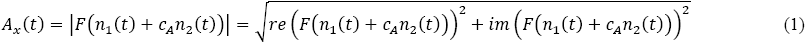

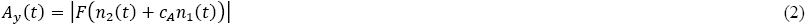

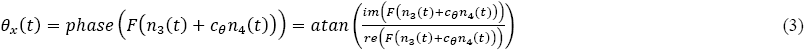

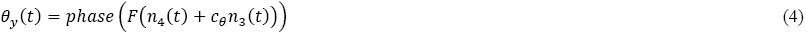

where *n*_*i*_ is a vector containing (*N*=50000) samples of Gaussian white noise from *i^th^* realization; *F* denotes complex Morlet wavelet transform with basis function *ψ*(*x*) = *e*^−*x*^2^/2^cos(5*x*); *c*_*A*_ and *c*_*θ*_ are scalar mixing parameters; *re* and *im* are the real and imaginary part of complex number, respectively; *A* and *θ* are the amplitudes and phases, respectively. This approach allows us to model phase and amplitude interactions separately [Bruns et al., 2000].

At the second stage, the amplitudes and phases (Eqs. 1–4) were used to assemble complex-valued time series in the following manner,

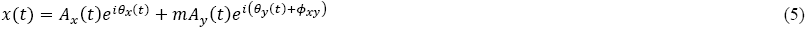

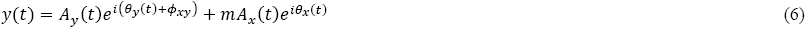

where *m* is the spatial mixing parameter, modelling the instantaneous signal spread; *φ_xy_* is the phase shift [-*π*, *π*], controlling the mean phase difference across sources x and y.

To demonstrate the spatial effects of signal spread, we simulated source signals in a 13×13 square grid layout, with inter-source distance *d_g_*. The signal spread was modelled as a truncated 2-dimensional Gaussian function with parameters *μ* = 0 and σ = *d_g_* up to a range of three standard deviations σ so that,

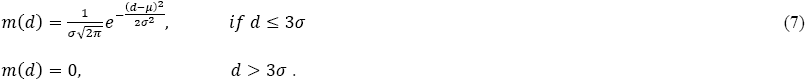

### 2.2 Quantification of interactions

Interactions between oscillatory neuronal signals can be measured in a variety of ways, which either rely on measuring some consistency of phase differences, correlation of amplitudes, or on a combination of both [Bastos and Schoffelen, 2016]. Here, we quantified interactions in terms of Phase Locking Value (*PLV*), and in terms of the correlation coefficient (*CC*) of amplitude envelopes. In addition, we used the imaginary part of the complex-valued PLV (*iPLV*) and the correlation coefficient of orthogonalized amplitude envelopes (*oCC*) to account for the effects of linear mixing.

Phase locking value (*PLV*) and imaginary part of phase locking value (*iPLV*) quantify the strength of phase coupling. *PLV* is defined as the magnitude of mean complex phase difference between amplitude-normalized source time courses [Lachaux et al., 1999],

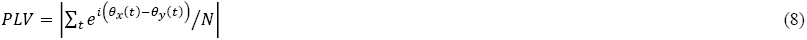

where *N* is the number of samples; |*·*| denotes absolute value operator; *θ_x_*(*t*) and *θ_y_*(*t*) are the phases of *x*(*t*) and y(*t*), respectively. *iPLV*, on the other hand, is the imaginary part of the average,

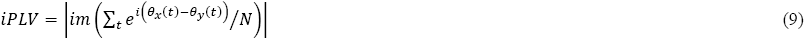

Thus, *PLV* theoretically compares to *iPLV* as coherence compares to the imaginary part of coherency [Nolte et al., 2004]. Nevertheless, it is important to keep in mind that the reliability of phase estimation inherently depends on SNR and may generally be more accurate in the presence of higher signal amplitudes [Palva et al., 2010]. Using the imaginary part, and thus discarding all real-valued contributions to the estimated interactions, effectively discards all zero-lag interactions, most of which are caused by instantaneous mixing and thus are considered detrimental to correlation estimates.

Amplitude correlations were quantified using the Pearson correlation coefficient (*CC*) between amplitude envelopes of *x*(*t*) and *y*(*t*), *A_x_*(*t*) and *A_y_*(*t*),

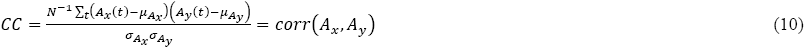

where *N* is the number of samples in signals *x*(*t*) and *y*(*t*); *μ_Ax_* and *σ_Ax_* refer to the average and standard deviation of *A_x_* over time, respectively.

Linear mixing between two signals *x*(*t*) and *y*(*t*) also affects the correlation between their amplitude envelopes. To exclude mixing-related amplitude correlations, two approaches [Brookes et al., 2012; [Hipp et al., 2012] have been proposed, where the signals are orthogonalized prior to the calculation of *CC*. This orthogonalization removes all linear contribution from signal *x(t)* to signal *y(t)*, or vice versa, provided that the signals are Gaussian—residual zero-lag mixing may remain for non-Gaussian signals [Brookes et al., 2014]. In the time domain, orthogonalization of signal *y* with respect to signal *x* is achieved as follows:

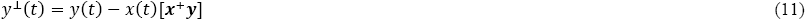

where ***x^+^*** is the pseudoinverse of the vector ***x*** [Brookes et al., 2012].

Alternatively, orthogonalization can be performed in frequency domain as follows [Hipp et al., 2012]:

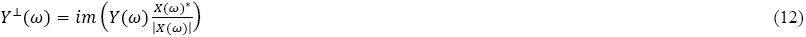

where^*^ denotes complex conjugation.

The orthogonalized *CC* (*oCC*) is then computed as *CC*, but using orthogonalized amplitude envelopes,

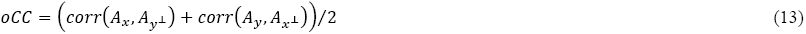

Because this seed-based orthogonalization can be performed in two directions, either to obtain *y^⊥^*(*t*) orthogonalized in relation to *x*(*t*), or to obtain *x^⊥^*(*t*) orthogonalized in relation to *y*(*t*), the final *oCC* is defined as the average of the two correlation coefficients. Such orthogonalization works, however, only under the assumption of data being normally distributed, which might not be accurate for the typically heavy-tailed oscillation amplitude distributions. It should also be noted that more sophisticated approaches for estimating amplitude-amplitude correlations, which simultaneously orthogonalize all the time series and greatly reduce spurious connections, have been introduced recently [Colclough et al., 2015; O’Neill et al., 2015]

In addition to *iPLV*, we also estimated the weighted phase lag index (*wPLI*) where the sign of the phase difference between two signals is weighted by the magnitude of the imaginary component of the cross-spectrum [Vinck et al., 2011],

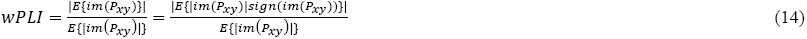

Where E{} is the expected value, im() is the imaginary part of a complex value, *P_xy_* is the cross-spectrum, *P*_*xy*_ = *x*(*t*)*y*^*^(*t*), x and y are complex signals, and ^*^ denotes the complex conjugate.

### 2.3 Simulations using realistic anatomical information and sensor topology

In addition to the synthetic simulations on the 2-dimensional ‘source’ grid, we investigated the effect of spurious synchrony in more realistic MEG/EEG settings. To this end, we simulated two correlated cortical parcels (left and right visual cortex) in a realistic anatomy and measurement geometry, and so that all other cortical parcels were given uncorrelated time series with equal amplitude distributions. The example parcels thus differed from others only by their correlation. We then performed a virtual MEG/EEG experiment by forward-modeling simulated source activity, followed by minimum-norm source reconstruction. Subsequently, we estimated all-to-all cortical interactions using the metrics outlined below.

#### Cortical reconstruction and parcellation

Volumetric segmentation of individual MRI images, reconstruction of anatomical surfaces, and cortical parcellation with Destrieux parcellation ([Dale et al., 1999; Fischl et al., 1999; Fischl et al., 2002]) were carried out with FreeSurfer (http://surfer.nmr.mgh.harvard.edu). The resulting parcellation contained 148 parcels covering the entire cortex. The largest parcels were iteratively selected and further partitioned until a total 400 parcels of equal size was obtained, see [Palva et al., 2010] for details.

#### Forward modeling

A realistic forward model was based on MRI data from one healthy subject (male, 32 years of age). T1-weighted anatomical MRI scans of were obtained at a resolution of 1×1×1 mm with a 1.5-T MRI scanner (Siemens, Germany). MNE-suite (*http://www.nmr.mgh.harvard.edu/martinos/userInfo/data/sofMNE.php*) was used to build a source model with 8196 current dipoles distributed evenly on the surface of the white matter and oriented normally to the local cortical surface. Also, a 3-layer MEG and EEG volume conduction model was created, which was used with the source model to construct the gain matrix *G*, using the linear collocation boundary-element method (BEM), as implemented in MNE-Suite. MEG and EEG sensor positions with respect to the head were taken from a concurrent MEG/EEG recording session [Palva et al., 2010].

#### Inverse modeling

We used *L^2^* minimum-norm estimation, as implemented in MNE Suite Matlab toolbox, to obtain a distributed cortical current estimate from the simulated sensor-level data. The inverse operator matrix *M* was computed as *M* = *RG*^T^(*GRG*^T^ + *λ*^2^*C*)^−1^, where regularization parameter *λ*^2^ = 0.1. The noise covariance matrix C was computed from empty-room noise for the MEG part; for the EEG part, an identity matrix was used (see for details, see [Palva et al., 2011]). The source covariance matrix R was set to an identity matrix.

#### Simulated cortical sources

We first simulated independent time series of 50,000 samples for each cortical parcel. We next simulated a ground truth interaction as correlation between two visual areas (Eqs. 1-4; see Fig. 6 for their anatomical locations). We then simulated EEG/MEG sensor data by forward-modeling these parcel time series to acquire sensor time series. Sensor time series were subsequently inverse-modeled to acquire reconstructed 8196 source time series, which was in turn collapsed into 400 parcels using a sparsely weighted collapse operator for optimal modeling accuracy [Korhonen et al., 2014]. Finally, we estimated all-to-all connectivity with *oCC, iPLV* and *wPLI.* For *oCC* estimation, we simulated coupling with *c_A_* = 0.9, *c_Θ_* = 0; for *iPLV* and *wPLI* estimation, we simulated coupling with *c_A_* = 0, *c_Θ_* = 0.9 and a phase difference of *φ*_xy_ = π/2.

The cortical spread of spurious correlations is determined by the cross-talk function (CTF), which describes how other sources influence the reconstructed time series of a source of interest. The CTF is obtained for the *n*-th cortical source as the *n*-th row of the product of the inverse and forward solutions, CTF(*n*) = (MG)*_n_* [Hauk et al., 2011], which we denoted as parcel-to-parcel *PLV_0_* (Fig. 6).

## 3 Results

To illustrate the concepts of artificial and spurious connections, we examine how variable linear signal mixing affects measures of phase and amplitude correlations under variable strengths of true phase and amplitude correlations. We aim here (1) to illustrate that spurious correlations which arise from linear mixing will be detected by interaction metrics supposed to be insensitive to linear-mixing, and (2) to characterize how the interpretation of phase and amplitude correlation measures is confounded by the interaction between linear mixing and the phase of true interactions.

### 3.1 Phase-locking value yields false positive correlations in the presence of signal mixing

Phase-locking value (*PLV*) is a commonly used measure of phase consistency between two time series [Lachaux et al., 1999]. *PLV*, like coherency and phase coherency in frequency domain, is sensitive to linear mixing of source signals in MEG and EEG recordings. A well-known example is that a single neuronal source (*e.g*. a cortical current dipole) generates strong and widespread channel-to-channel correlations [Schoffelen and J. Gross, 2009]. Figure 1A-C illustrates the effect of signal mixing on the PLV. We first simulated two signals that were phase-coupled with a phase lag of 54 deg (norm. phase lag of 0.3, see Fig. 1A, *m* = 0) and quantified their phase difference distribution, which as expected peaks at the simulated phase lag (Fig. 1B, green). Linear mixing of these time series (mixing parameter *m* = 0.4, see Eq. 5, Fig. 1A *bottom half*) has two effects on the phase difference distribution: it becomes narrower, *i.e.*, phase difference between *x* and *y* was observed as being more consistent, and the peak is shifted towards zero (Fig. 1B). These effects are reflected in the changes in the magnitude and phase, respectively, of complex-valued average phase difference vectors (Fig. 1C).

**Fig. 1.**
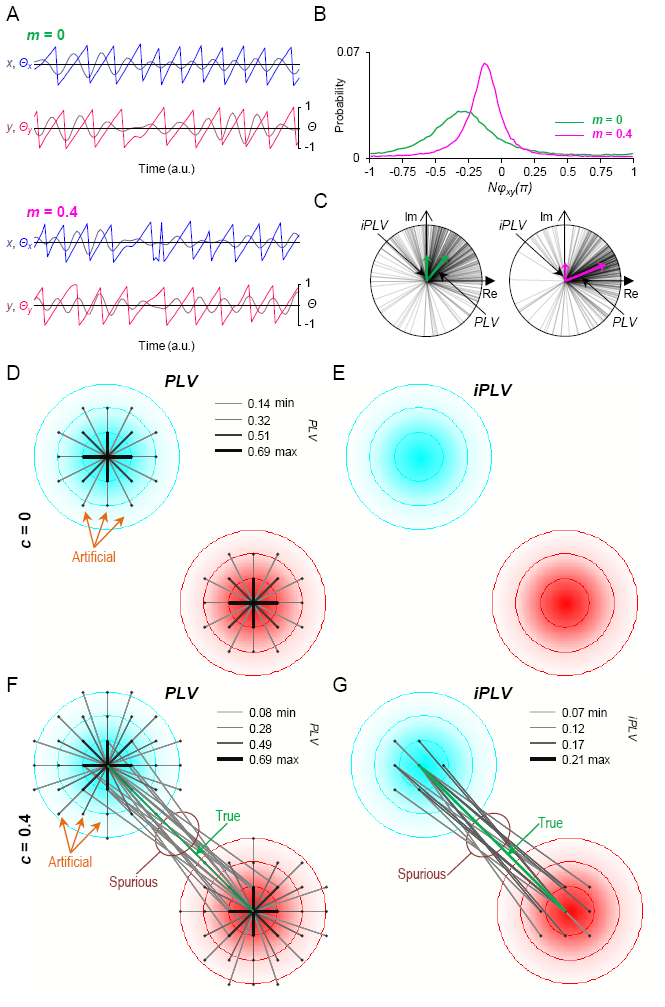
*PLV* and *iPLV* measure the strength of phase correlation but are biased by signal mixing. A) Coupled (*c* = 0.4) real-valued signals *x*(*t*) and *y*(*t*) and their phases *Θ_x_*(*t*) and *Θ_y_*(*t*) in the absence (*m* = 0) and presence (*m* = 0.4) of linear mixing. B) Distribution of the phase difference *ø_xy_* with (*m* = 0.4) and without (*m* = 0) linear mixing. The true phase difference (*ø_xy_*) = −0.3π. C) Vector interpretation of the distributions in B. Left: without mixing, right: with mixing. Increasing linear mixing biases phase difference distribution towards *ø_xy_* = 0, therefore increasing *PLV* while decreasing *iPLV*. D) Mixing causes false positive *artificial PLV* interactions, *within* the mixing region even in the absence of true correlations. Activity of 169 uncoupled (*c* = 0) sources (black dots) placed into a 13x13 grid was simulated and the 20 strongest *PLV* edges of the two sources-of-interest (centers of the cyan and red regions) were picked for visualization. The cyan and red color gradients indicate mixing strength. No supra-threshold *PLV*s occur between sources that are not linearly mixed. E) *iPLV* analysis of the same data as in D shows that *iPLV* does not discover artificial interactions. F) True phase correlations are mirrored into false positive *spurious* correlations, between *different* mixing regions when there is a true interaction (*c* = 0.9) between two sources-of-interest (centers of the mixing regions). Note that the strongest edges detected were artificial. G) *iPLV* does not discover *artificial* interactions, but it detects spurious interactions similarly to *PLV*. Spurious correlations arise because any two sources in separate mixing regions partially retain the non-zero phase difference of their center sources.

Figs 1D-G illustrate the effect of signal spread on the estimated interactions, and illustrate the distinction between what we defined as artificial and spurious interactions. We simulated source reconstructed data on a 13x13 grid with a well-defined point-spread characteristic, as defined in equation 7, and computed all pairwise interactions between the reconstructed sources. The cyan and red contours in figs 1D-G specify the point-spread for the two sources at the centre of these regions. The grayscale of the edges connecting source locations reflect the estimated interaction strength between the reconstructed signals, after signal mixing. Prior to mixing, the activity of two of the sources (the central nodes of the cyan and red regions in Figs 1D-G) was coupled by non-zero phase lag with a coupling strength *c*. Figs. 1D-E show the estimated *PLV* and *iPLV* when *c* was set to 0, *i.e*. no phase correlations. After source mixing, the *PLV* (Fig. 1D) shows strong local artificial interactions, which are not visible in the *iPLV* (Fig. 1E). These false positive, *artificial* connections are caused directly by signal mixing and have unimodal phase difference distributions centered around zero-lag.

Figs. 1F-G show a simulation where a true phase correlation was introduced between the central sources of the cyan and red regions (*c* = 0.4). This true coupling still resulted in local artificial interactions in the *PLV* (Fig. 1F), which were abolished in the *iPLV* (Fig. 1G). Importantly, apart from revealing the true interaction (green lines in 1F and 1G), many spurious interactions were present, both using *PLV* and *iPLV* as interaction measure.

### 3.2 Linear-mixing insensitive phase-locking measures do not eliminate spurious correlations

The imaginary part of the complex *PLV* (*iPLV*, Fig. 1C) also indexes phase consistency of the two time series but like its frequency-domain homolog, imaginary coherency, it is insensitive to the direct effects of linear mixing that have zero-phase-lag and are reflected in the real part of the complex interaction metric [Nolte et al., 2004; Vinck et al., 2011]. The insensitivity of *iPLV* to instantaneous linear mixing is clear in the grid-source simulation where in the absence of true phase-lagged coupling, no significant correlations were detected (Fig. 1E). In the presence of a true phase correlation (as in Fig. 1F), this correlation was correctly identified by *iPLV* (Fig. 1G). However, like *PLV*, *iPLV* also discovered dense spurious correlations, *i.e.*, ghost interactions in the vicinity of the true connection. Thus, even if *iPLV* correctly rejects within-region signal mixing effects, it is as sensitive to spurious correlations as *PLV* is.

Taken together, in the presence of signal spread, any bi-variate measure that estimates phase coupling influenced by linear mixing will yield both artificial and spurious false positive observations, whereas measures insensitive to instantaneous mixing do not detect the artificial correlations but they do yield spurious interactions, *i.e.*, the ghost edges surrounding true interactions (Fig 1 G).

### 3.3 Correlation coefficient produces artificial and spurious amplitude correlations

Figure 2 demonstrates the effect of signal spread on amplitude correlation measures. We simulated two amplitude-coupled signals and computed the correlation coefficient (*CC*) between the signals’ amplitude envelopes before and after signal mixing at *m* = 0.4. As expected, signal mixing increases the similarity between amplitude envelopes (Fig. 2A) and strengthens *CC* (Fig. 2B).

**Fig. 2.**
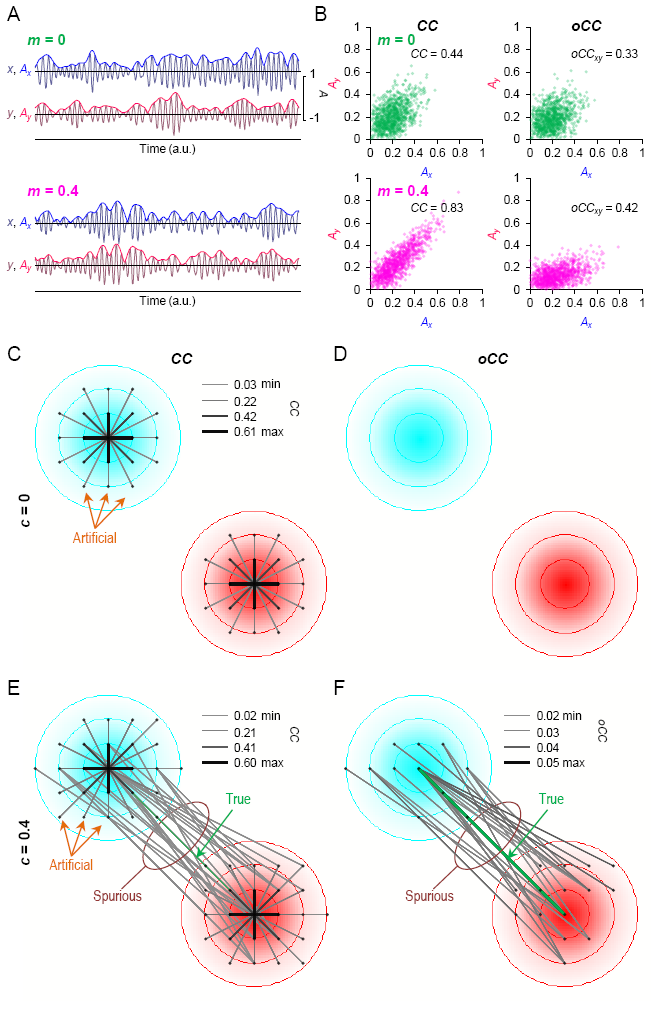
Measures of amplitude correlation, *CC* and *oCC*, are corrupted by signal mixing similarly to estimates of phase correlation. A) Coupled (*c* = 0.4) real-valued signals *x*(*t*) and *y*(*t*) and their amplitude envelopes *A_x_*(*t*) and *A_y_*(*t*) in the absence (*m* = 0) and presence (*m* = 0.4) of linear signal mixing. The true amplitude correlation is artificially amplified by the linear mixing. B) *A_x_*(*t*) and *A_y_*(*t*) values with and without linear mixing for estimation of *CC* (left; each dot represents a sample) and *A_x_*(*t*) and orthogonalized *A_y_*(*t*) values for estimation of *oCC*. C) *CC* is biased by linear mixing similarly as *PLV* (visualization and simulations as in Fig. 1 D) D) *oCC* is insensitive to artificial correlations similarly to *iPLV* (data as in C). E) True correlation interaction is surrounded by spurious edges in *CC* interaction. The strongest edges detected were artificial. F) *oCC* ignored the artificial correlations, as in D. However, orthogonalization did not solve the problem of spurious edges: *oCC* detects spurious correlations similarly to *CC* (data as in E).

In the source-grid analysis, when true correlations were not present, the mixing of random source signals produced region-constrained artificial amplitude correlations (Fig. 2C), exactly as found for *PLV* (see Fig. 1D). In the same vein, a true correlation was accompanied by long-range spurious *CC* between the coupled regions, in addition to the artificial correlations (Fig. 2E). Hence *CC*, similarly to *PLV*, yields both artificial and spurious observations of amplitude correlations in the presence of signal mixing.

### 3.4 Orthogonalized correlation coefficient produces spurious amplitude correlations

Orthogonalization, *i.e.*, the removal of linear dependencies, of the two real-valued signals, *x*(*t*) and *y*(*t*), before the estimation of the amplitude envelopes and their correlation, excludes the contribution of linear mixing to the correlation estimates [Brookes et al., 2012; [Hipp et al., 2012].

Similarly to *CC*, *orthogonalized CC (oCC)* identifies the correlation between two coupled simulated signals. After linear mixing and orthogonalization of signal *y* with respect to *x*, the *oCC* between *A*_y_ and *A*_x_ was smaller than *CC*, but still greater than the *oCC* obtained before mixing (Fig. 2B).

The insensitivity of *oCC* to artificial amplitude correlations was clear in grid-model simulations. A mixing of random source time courses did not lead to significant *oCC* between any sources (Fig. 2D). However, when a true amplitude correlation was present, it was mirrored into multiple FP spurious correlations in estimated *oCC* interaction matrix, which are shown as widespread ghost edges in the synchrony graph (Fig. 2F). Thus, amplitude correlations estimated with *oCC* share the caveats of phase correlations identified with *iPLV*.

### 3.5 *PLV* and *iPLV* are differentially sensitive to signal mixing and phase difference

Next we assessed the effect of linear mixing on the *PLV* and *iPLV* estimates under different regimes of true phase coupling and phase differences (Fig. 3).. We simulated two signals *x*(*t*) and *y*(*t*) and parametrically varied their phase coupling (*c_Θ_* = 0 … 1; Eqs. 3–4), phase difference (*ø_xy_* = −π … π) and linear mixing (*m* = 0 … 0.6; Eqs. 5–6). For each combination of these parameters, we computed the *PLV* and *iPLV*. Fig. 3A shows the effect of the amount of actual phase coupling on the estimated *PLV* under various amounts of linear mixing, keeping the phase difference fixed at 0.3 (norm. phase). Fig. 3C shows the estimated *PLV* as a function of the phase difference, given a fixed amount of actual phase coupling of 0.4. Both panels show a strong nonlinear dependency of the coupling strength and the phase difference on the estimated *PLV*, which in itself depends on the amount of linear mixing. At low phase differences in particular, the *PLV* shows a positive bias, which increases with the strength of signal mixing. This observation can be explained by the fact that the relative contribution of the zero phase lag linear mixing to the estimated *PLV* works ‘synergetically’ with the true coupling at small phase differences, whereas it has a ‘counteracting’ effect when the phase difference of the true coupling is far away from 0. Moreover, this effect saturates at higher values of true coupling, because the *PLV* is by definition bounded to a maximum value of 1. For the same set of simulations, Fig. 3B and 3D show the *iPLV*. In contrast to *PLV*, linear mixing reduces the estimated *iPLV* for all *c_Θ_*, at a fixed phase difference of 0.3π, and most strongly does so for large *c_Θ_* values (Fig. 3B). *iPLV* is reduced by increasing signal mixing because the phase difference distribution shifts towards zero with increasing mixing (Fig. 1B).

**Fig. 3.**
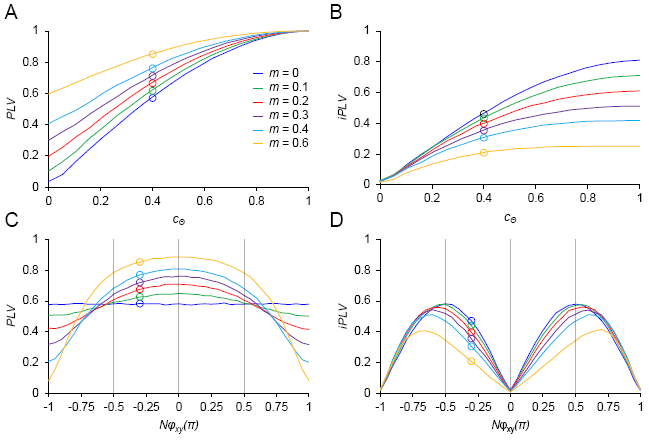
*PLV* and *iPLV* are affected by phase coupling strength *c_Θ_*, phase difference *nϕ_xy_* and linear mixing *m*. Phase coupling and linear signal mixing (*m* = 0 (blue), 0.1 (green), 0.2 (red), 0.3 (violet), 0.4 (cyan), and 0.6 (orange)) were simulated between two signals. A) *PLV* between the signals as a function of *c_Θ_* and *m*, when *nø_xy_* = −0.3. Open circles at *c_Θ_* = 0.4 visualize the coupling strength used in C and D. B) *iPLV* between the signals as a function of *c_Θ_* and *m*. C) *PLV* as a function of *nø_xy_*, when *c_Θ_* was set to 0.4. PLV is greatly affected by the phase difference when signal mixing is strong. Open circles at *nϕ_xy_* = −0.3 visualize the *nϕ_xy_* used in A and B. D) The strength of *iPLV* depends highly on the phase difference, and it is biased towards large phase difference; *iPLV* is abolished when *nϕ_xy_* = 0 or *nϕ_xy_* = ±p.

Algebraically, *PLV* (Eq. 8) is independent of the mean phase difference, *φ_xy_*. Yet, in the presence of linear mixing, the estimated *PLV* is dependent on *φ_xy_* (Fig. 3C). The bias from *φ_xy_* on the estimated *PLV* can be positive or negative: it is positive for small phase differences (*φ_xy_* < π/2, i.e. 90 deg) because under such a regime the interaction and mixing effects add up. Consequently, the bias is negative for near “anti-phase” narrow-band synchrony when *φ_xy_* approaches ±π, and *x*(*t*) and *y*(*t*) have reverse polarities. The case is, again, very different for *iPLV*, which is zero for *ϕ_xy_* = 0 and ±p, as can be seen from its definition (Eq. 9). Between these poles, *iPLV* is always negatively biased by signal mixing, regardless of the mean phase difference (Fig. 3D).

Taken together, *iPLV* is not positively biased by signal mixing as *PLV* is, although it has the disadvantage of failing to detect true synchronizations that are near zero‐ or anti-phase-lag. The properties of these two phase correlation measures lead to an interesting worst-case scenario, where phase synchrony is accompanied by strong signal mixing and *iPLV* = 0. Then *PLV* can have almost any value, depending on the actual *c_Θ_* and whether *φ_xy_* = 0 or ±π.

### 3.6 *CC* and *oCC* are biased by signal mixing and phase effects

In our grid simulations, the behavior of *CC* and *oCC* was almost identical to that of *PLV* and *iPLV*, respectively. We asked next if their similarities extend to phase effects. Phase effects are not often considered in studies of amplitude correlations, because *CC* and *oCC* are thought to quantify the correlation between amplitude envelopes and amplitude is independent of phase.

We first simulated two signals *x*(*t*) and *y*(*t*) and parametrically varied their amplitude coupling (*c_A_* = 0 … 1; Eqs. 1–2), phase difference (*ø_xy_* = −π … π) and linear mixing (*m* = 0 … 0.6; Eqs. 5–6). We then computed *CC* and *oCC* for each combination of parameters. In the absence of any concurrent phase correlations, signal mixing introduced as expected a positive bias on the *CC*, in particular at low to intermediate values of *c*_A_ (Fig. 4A). The effect of signal mixing was different for *oCC*, in close resemblance to what was found for *iPLV* (Fig. 3B): signal mixing drastically reduced *oCC* for high values of *c*_A_ (Fig. 4B).

**Fig. 4.**
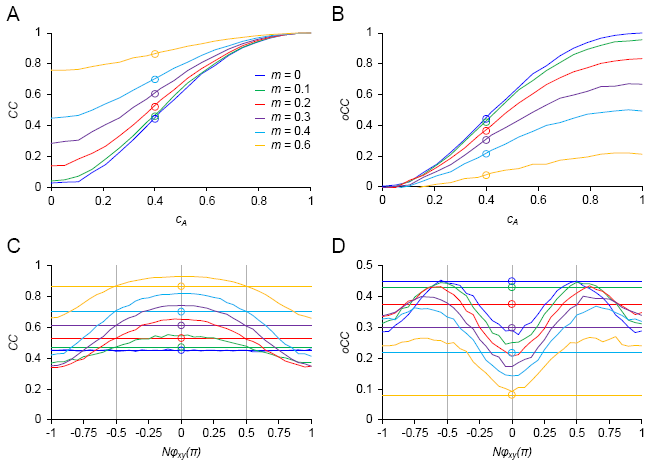
*CC* and *oCC* (Brookes et al, 2012) vary as a function of amplitude coupling strength *c_A_*, phase difference *nϕ_xy_* and linear mixing *m*. Two signals with distinct coupling strength for amplitude and phase (*c_Θ_*) were simulated and linearly mixed. A) *CC* between the signals as a function of *c_A_* and *m* in the absence of phase correlations (*c_Θ_* = 0, *nϕ_xy_* = 0). Open circles at *c_A_* = 0.4 visualize the coupling strength used in C and D. B) *oCC* between the signals as a function of *c_A_* and *m*. C) *CC* between the signals as a function of *nϕ_xy_* of the phase coupling, when *c_Θ_* = 0.4. The horizontal lines visualize the mean *CC* obtained at *c_Θ_* = 0. Open circles mark *nϕ_xy_* = 0 and *c_Θ_* = 0 used in A and B. D) *oCC* between the signals as a function of *nϕ_xy_*, when *c_Θ_* = 0.4. Note that when a phase coupling is present in addition to the amplitude coupling, both *CC* and *oCC* are biased by *nϕ_xy_*, but in different manners.

Interestingly, introduction of a true phase correlation (*c_Θ_* = 0.4) resulted in a phase different dependent effect on the estimated *CC* and *oCC*, with a variable influence of signal mixing (Fig. 4C and 4D). Comparing the straight lines (representing the absence of phase coupling) with the curves of the same colour (representing the presence of phase coupling), signal mixing increased the estimated *CC* when the phase difference *φ_xy_* was small and reduced the estimated *CC* when the signals were close to anti-phase, *i.e*. *ø_xy_* = ±π (Fig. 4C). This is because the phase correlation at small phase differences leads to an alignment of the peaks of *x*(*t*) and *y*(*t*). Now, if *A*_x_ and *A*_y_ are in fact correlated, the linear mixing effectively ‘amplifies’ this correlation, because high peaks will match with high peaks more often than with low peaks (and low peaks will match more often with low peaks than with high peaks).

For the estimated *oCC*, the presence of actual phase coupling affected the estimates in a nonlinear and phase difference dependent way. This is a result of the orthogonalization process (see Eqs. 11–12), which, before computing the amplitude correlation, implicitly either regresses out the real valued contribution of signal *x*(*t*) to *y*(*t*) (Eq.11) or explicitly only uses the imaginary component of the cross-terms between signals *x* and *y* (after amplitude normalization for one the signals). Either way, the real/imaginary part of a complex-valued signal mixes phase information with amplitude information. A deviation from a uniform distribution of phase differences across observations (*i.e.*, the presence of phase coupling), will affect the orthogonalization process in a non-trivial way, despite the fact that consecutively only the amplitude terms are used to compute the correlation. Our simulations show that phase correlations do indeed have an impact on *oCC*. In the presence of phase correlations and linear mixing, the estimated *oCC* is reduced when the mean phase difference is close to 0 (Fig. 4D). On the other hand the estimated *oCC* is inflated when *ϕ_xy_* is close to ±π/2. These phenomena can be understood from the properties of orthogonalization, where the orthogonalized amplitudes of a signal are obtained by projecting the complex-valued phase-amplitude vector onto the imaginary axis, after rotation with the phase of the other signal. A consistent phase relationship across observations (phase coupling) will amplify the estimated correlation due to a consistent rotation of the single observation phase-amplitude vectors towards the imaginary axis, thus increasing the contribution of the spatially leaked amplitudes. A consistent phase relationship of around 0 will result in a consistent absence of vector rotation prior to imaginary axis projection, and any spatially leaked amplitude components will be lost when taking the imaginary component. Phrased differently, for highly similar time series, the resulting orthogonalized signal will be almost negligible, *i.e*. *y^⊥^*(*t*) ≈ 0, leading to small envelope correlation values. On the other hand, when the phase difference between *x*(*t*) and *y*(*t*) are mostly at ±π/2, they are considered already orthogonal and are barely affected by the orthogonalization procedure, *i.e*. *y^⊥^*(*t*) ≈ *y*(*t*), even if there are correlations induced by signal mixing.

Hence, for a range of values of phase lags, *φ_xy_*, the estimated amplitude correlation can be significantly affected by the presence of concurrent phase coupling. To get a more complete picture of the interaction between *c_A_*, *c_Θ_* and *ø_xy_*, we extended these simulations for a large part of the parameter space and for both the regression‐ and the imaginary-projection-based orthogonalization methods (Supplementary Figs S1 and S2). Both methods were approximately equally affected by true neuronal phase correlations, both in the presence and in the absence of linear mixing. These findings thus show that *oCC* produces false positive amplitude-correlation observations in the absence of any true amplitude correlations when true phase correlations are present (see Figs S1-S2 for *c_A_* = 0 and *c_Θ_* is high).

### 3.7 Weighted phase-lag index (wPLI) estimates of phase coupling are not biased by mixing

The *wPLI* estimates the extent to which phase leads and lags between two signals are non-equiprobable, and it weighs the observations by the magnitude of the imaginary component of the cross-spectrum [Vinck et al., 2011]. Unlike what was observed with *iPLV*, linear mixing does not affect the *wPLI* estimates across the tested range of coupling strengths (Fig 5A) or over different phase differences of a true correlation, *c_Θ_* = 0.4 (Fig 5B). Taken together, *wPLI* estimates are not affected by mixing as *iPLV* estimates are, but are still compromised in overall utility by the phase-difference dependence of the metric value and its inability to detect true near zero‐ or anti-phase-lag phase synchronizations. Moreover, wPLI is only insensitive to mixing for not more than two sources.

**Fig. 5.**
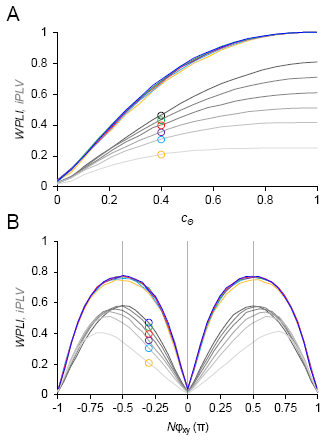
A) Linear mixing positively bias *wPLI* estimates, but the amount of mixing does not seem to differentiate *wPLI* estimates like what was observed with *iPLV* (gray lines, cf Fig. 3B). B) When a true phase interaction is present (*c_Θ_* = 0.4), mixing does not bias *wPLI* estimates at any of the tested phase lags (*n*ϕ_xy_)

### 3.8 All linear-mixing insensitive interaction metrics produce wide-spread spurious synchrony in a realistic simulations

To demonstrate the effect of spurious synchrony in real MEG/EEG settings, we performed a simulation using a realistic model based on individual MR images and a real MEG/EEG measurement geometry. We simulated independent time series across the whole cortex except for two highly correlated sources located in left and right visual areas. After a virtual MEG/EEG experiment, *i.e.*, forward‐ and inverse-modeling of the simulated time series, we estimated all-to-all connectivity and visualized the extent of spurious phase correlations in synchrony graphs displayed together with the magnitude of cross talk on a flattened cortical maps. (right column, Fig. 6). All tested interaction metrics, iPLV, wPLI, and oCC yielded significant amounts of ghost connections (grey) around the true connection (black) among parcels of which the signals were mixed with those of the two truly connected parcels. Cross talk was measured here among all parcels by *PLV* estimates of forward‐ and inverse-modeled filtered noise parcel signals. To provide another example with more distant true sources than the two visual ones, we performed a similar analysis where a true interaction was simulated between middle frontal gyrus and inferior parietal gyrus (Fig. S3A). This analysis revealed a qualitatively identical result with ghost/spurious connections surrounding the true connection. These realistic simulations thus show that ghost connectivity is a tangible problem in MEG source space connectivity analyses and involves significant distances across the cortical surface. Importantly, the problem cannot be alleviated by picking a coarse parcellation resolution as adjacent parcels will express mixing in any case. To illustrate this, we displayed the parcel-parcel crosstalk of a parietal and frontal parcel with their surroundings for the Desikan-Killiany atlas (68 parcels), the Destrieux atlas and its subdivisions (148, 200, 400 parcels), and the source dipoles *per se* of the cortical source model (6400 sources, Fig. S3B). Significant mixing of similar spatial extent was visible at all resolutions.

**Fig. 6.**
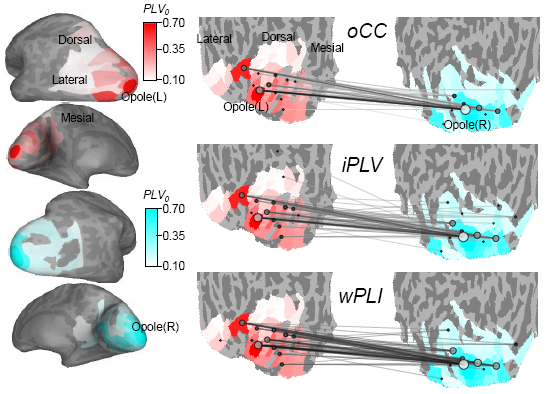
Left: Illustration of the mixing effect, quantified with parcel-to-parcel *PLV_0_*, simulated parcel time series data on a 3D model of brain of one subject. The colour gradient on the flattened cortical map indicates the intensity of mixing from the simulated parcels. Red: mixing of left occipital pole (Opole). Cyan: right Opole. Right: amplitude and phase coupling were simulated between left and right Opole while the rest cortical parcels time series was uncorrelated. Simulated time series were forward‐ and inverse-modeled and estimated with *oCC*, *iPLV* and *wPLI.* The strongest 60 edges were overlaid on flattened cortical map. oCC graph (Brookes, 2012) was computed using time series that was simulated with *c_A_* = 0.9, *c_Θ_* = 0. *iPLV* and *wPLI* graphs were computed using time series that were simulated with *c_A_* = 0, *c_Θ_* = 0.9, *nϕ_xy_* = −0.5.

## 4 Discussion

In recent years, connectivity measures that ignore zero-phase-lag interactions have been developed to protect interaction estimates against inflation and false positive findings by linear mixing of the underlying signals, which is an unavoidable phenomenon in MEG and EEG research. In this study, we question the often written claim that, in the presence of true interactions, such coupling measures (such as *iPLV*, *wPLI* and *oCC*) would be *de facto* immune to false positive detections. Although these measures can be overly conservative by missing true near-zero-phase interactions, we show here that they also yield false positive interactions due to signal spread. This is because field spread in the vicinity of a true non-zero phase interaction gives rise to spurious “ghost” interactions, that appear as false positives with any bivariate interaction measures. Moreover, indicating further interpretational challenges, our simulations showed that orthogonalized amplitude correlation coefficients are not independent of concurrent phase coupling. In fact, they are non‐trivially affected by the presence of true phase coupling and linear mixing in a phase-difference dependent manner and may yield both false positive and negative findings.

Our simulations illustrate the expected effect of volume conduction or field spread on standard measures of amplitude-amplitude and phase-phase coupling: in the presence of linear mixing, *CC* and *PLV* estimates yield artificially inflated coupling estimates for sources with a true interaction. Notably, this phenomenon leads to purely artificial coupling even when two source time series are uncorrelated. Our simulations also corroborated earlier studies by showing that modified versions of these measures that are insensitive to instantaneous coupling (*i.e.*, *oCC* and *iPLV/wPLI*) detected no coupling in the absence of a true interactions despite of the presence of signal spread. However, in the presence of a true interaction, signal spread produces spurious “ghost” interactions among uncorrelated sources in the vicinity of the truly interacting sources.

Furthermore, we show that signal spread affects different interaction estimates in different ways. Notably, the presence of linear mixing leads to an inflation of the estimated *PLV* and *CC*, but to an underestimation of true coupling when using *iPLV* or *oCC*. This is an important observation as it challenges the widely supported claim that the interpretation of signal spread insensitive coupling measures are not affected by linear mixing, although *wPLI* constitutes an important exception from this.

Moreover, we show that impact of signal mixing on both *PLV* and *iPLV* estimates is dependent on the phase difference of the true coupling. Hence changes in phase differences in the absence of changes in coupling strength between contrasted conditions would appear as false positive changes in coupling strength in a signal-mixing dependent manner. Moreover, while *wPLI* is independent of signal mixing here, it is still very dependent on the phase difference and would thus be similarly confounded.

As a major methodological finding, we observed that phase coupling and its phase difference influenced also the *CC* and *oCC* estimates of amplitude correlations among oscillations. These latter results indicate that phase coupling among the signals can impact the estimation of amplitude coupling, and this effect is amplified with increasing amounts of linear mixing. This poses serious limitations on distinguishing pure phase‐ from pure amplitude-coupling phenomena, and more generally limits the interpretability of such measures in isolation.

In summary, our simulations, including realistic MEG/EEG configuration, illustrate two main problems that need to be acknowledged to avoid false interpretations of connectivity analyses. First, we show that using measures that do not detect zero-phase coupling is by no means a guarantee against false positives. As acknowledged, these measures are indeed not affected by artificial coupling caused by linear mixing, but they are still prone to detecting ghost connections, *i.e.*, spurious interactions that arise in the vicinity of true interactions. These “2^nd^ order false positives” are caused by the unavoidable cross-talk between the source estimates that is preserved at all resolutions of source parcellations. It is important to note that the spatial structure in the cross-talk function is generally not smooth as a function of distance to the source of interest. As a consequence, spurious interactions may arise at locations further away from the primarily interacting sources. The exact form of the cross-talk function is also a property of a specific inverse solution, but it universally leads to linear mixing among numbers of sources and thus the problem of spurious interactions is qualitatively identical to all source reconstruction approaches. Second, by showing the effects of phase correlations on amplitude correlation measures with varying amounts of linear mixing, we demonstrated the limitations on the separability of phase and amplitude interactions. In a worst-case scenario, for instance, linear mixing and strong phase coupling at around p/2 phase lag will lead to large values of the estimated *oCC* in complete absence of true amplitude correlations. This represents an extreme case of false positives that this measure can produce. Conversely, through the effect of signal-to-noise ratio on the accuracy of phase estimates, also phase correlations can be affected by amplitude dynamics and correlations [Palva et al., 2010].

We consider the above limitations to be of major importance to the EEG and MEG field. Our results confirm the added value of recently proposed coupling measures which focus on the non-zero phase interactions. However, they also reveal a number of limitations that have been either underrated or simply hardly taken into consideration. Most importantly, the red flag raised here based on simulations is valid for real data situations. The behavior of the coupling estimators was investigated by modulating all principal parameters that affect the measures used. Although we did not test the effect of additive noise, our main findings are expected to remain identical in the presence of noise. In addition, the reported limitations hold for both spontaneous and evoked data, with and without a contrast condition comparison. Moreover, all forms of cross-frequency or other non-linear couplings, albeit immune to the artificial interaction effect *per se*, will also equally suffer from the spurious/ghost coupling effect.

Since the main limitation of the *oCC* and *iPLV* methods shown here is that these measures also will yield ghost connections, mostly in the vicinity of the true connections, one might argue that the problem could probably be addressed by local selection of the edges with the highest coupling strength, using a clustering approach, or accept and exploit their presence by spatially non-homogeneous smoothing [Schoffelen and J. Gross, 2011]. With respect to the local selection of the strongest edge, it should be noted that the strongest or statistically most significant interactions will not necessarily correspond to the true interaction. A simple theoretical example would be a situation where there are two pairs of truly (and similarly) interacting sources. Source estimates at locations in between the individual nodes of each interacting source pair will be affected by cross-talk from the interacting nodes, which leads to spurious connections, of which the amplitude and statistical robustness may exceed that of the true interactions around it. Generally speaking, there is no guarantee that true interactions will exhibit a greater coupling strength or display higher statistical significance than ghost connections.

For this account to be forward-thinking and constructive, it is important to explore potential recommendations and suggestions that arise from our observations. The first recommendation is that one should understand and acknowledge the limitations of the source reconstruction and coupling method used when reporting the MEG/EEG connectivity results. Claims about ruling out false positives using methods insensitive to instantaneous coupling should be avoided. Likewise, it is essential to move from analyses limited to a few regions-of-interest into full source-space interaction mapping to avoid neuroanatomical misinterpretations of the coupled sources.. Restriction to a seed-based approach might imply that one might focus interpretations on a detected interaction, but will fail to notice potential coupling that exists in its vicinity or mistake a ghost interaction for a true one. Neighboring connections might in theory contain the true interacting pair of sources while the one revealed in a seed-based approach could simply be a ghost of the real interaction. Additionally, a general recommendation would be to also explore phase coupling even when the main interest lies in assessing amplitude coupling. We have shown that if strong phase correlation is present, linear mixing can lead to erroneous amplitude correlation estimations. Systematically assessing phase and amplitude coupling might therefore be very helpful when interpreting the findings.

Ultimately, finding the ideal measure to characterize interactions using MEG or EEG is limited by our knowledge of the true mechanisms of neuronal interactions. The best we can do is to estimate brain interactions with one or several methods for which we have a thorough understanding of the strengths and drawbacks. The limitations of the connectivity measures we choose to use need to be explicitly acknowledged and potential implications on the interpretation of the data need to be discussed. There are also new analysis possibilities, such as using multivariate correction [Brookes et al., 2014; Colclough et al., 2015; Soto et al., 2016] and hyper-edge bundling (Wang et al., submitted) approaches, that alleviate the problem of ghost interactions but each with their limitations. Beyond sounding the alarm, the current study intends to help improve good practice in MEG & EEG source connectivity analyses by outlining potential interpretational pitfalls and promoting some standards of good practice.

## Acknowledgments

This study was supported by the Academy of Finland Grants 253130 and 256472 to J.M.P. and 1126967 to S.P., and by the Helsinki University Research Funds. MJB was funded by a Medical Research Council New Investigator Research Grant (MR/M006301/1). KJ was supported by funding from the Canada Research Chairs program and a Discovery Grant (RGPIN-2015-04854) awarded by the Natural Sciences and Engineering Research Council of Canada.

## Figure legends

**Table 1** Division of interaction metrics into four groups by the correlation they measure (phase or amplitude) and their sensitivity to zero-phase lag interactions. (*cf*. Vinck et al., 2011)

**Fig. S1** When phase coupling is present, *oCC* is biased by the phase difference. To comprehensively visualize the dependency of *oCC* (as computed in {{1725 Brookes 2012;}}; Eq. 11) on *nϕ_xy_*, constant values were set for *c_A_* (*c_A_* = 0, *c_A_* = 0.2, and *c_A_* = 0.4), *c_Θ_* (*c_Θ_* = 0, *c_Θ_* = 0.1, *c_Θ_* = 0.2, *c_Θ_* = 0.3, *c_Θ_* = 0.4, *c_Θ_* = 0.5, and *c_Θ_* = 0.6) and *m*, and *nϕ_xy_* was varied between −1 and 1. When phase correlations are present, oCC is biased by *nϕ_xy_*, independent of linear mixing. Strong phase correlation (*c_Θ_* ≥ 0.4) leads to false positive *oCC* in the presence of linear mixing, even in absence of true amplitude correlations. Horizontal lines that show the mean *oCC* without bias from phase correlations (*c_Θ_* = 0) have been calculated as an average over the *nϕ_xy_* range with *c_Θ_* = 0 and *c_A_* = 0, *c_A_* = 0.2, and *c_A_* = 0.4 (the uppermost row).

**Fig. S2** Same as in Fig. S1, but *oCC* was calculated as described in {{1536 Hipp 2012;}} (Eq. 13). The two orthogonalization methods differed mostly in the strength of the mixing effect, when real amplitude correlations were present. In terms of *nϕ*_xy_ bias, there is no difference between the two methods.

**Fig. S3** A) Illustration of the mixing effect comparable to that in Fig. 6 but for a case where the true interaction was simulated between medial frontal gyrus (red) and the inferior parietal gyrus (cyan). B) Illustration of parcel-to-parcel mixing (as in A and Fig. 6) for multiple resolutions of cortical parcellations.

